# Pushing Raman spectroscopy over the edge: purported signatures of organic molecules in fossils are instrumental artefacts

**DOI:** 10.1101/2020.11.09.375212

**Authors:** Julien Alleon, Gilles Montagnac, Bruno Reynard, Thibault Brulé, Mathieu Thoury, Pierre Gueriau

**Affiliations:** Institute of Earth Sciences, University of Lausanne, Géopolis, CH-1015 Lausanne, Switzerland; Université de Lyon, ENS de Lyon, Université Lyon 1, CNRS, LGL-TPE, F-69007 Lyon, France; HORIBA France SAS, 91120 Palaiseau, France; Université Paris-Saclay, CNRS, ministère de la Culture, UVSQ, MNHN, Institut photonique d’analyse non-destructive européen des matériaux anciens, 91192, Saint-Aubin, France

**Keywords:** Raman, fossil biomolecules, biosignatures, Wavelet transform, Baseline subtraction, edge filter ripples

## Abstract

Claims for the widespread preservation of fossilized biomolecules in many fossil animals have recently been reported in six studies, based on Raman microspectroscopy. Here, we show that the putative Raman signatures of organic compounds in these fossils are actually instrumental artefacts resulting from intense background luminescence. Raman spectroscopy relies upon the detection of photons scattered inelastically by matter as a result of its interaction with a laser beam. For many natural materials, this interaction also generates a luminescence signal that is often orders of magnitude more intense than the light produced by Raman scattering. Such luminescence, coupled with the transmission properties of the spectrometer, induced quasi-periodic ripples in the measured spectra that have been incorrectly interpreted as Raman signatures of organic molecules. Although several analytical strategies have been developed to overcome this common issue, Raman microspectroscopy as used in the studies questioned here cannot be used to identify fossil biomolecules.

## Introduction

The fossil record contains unique information to document the evolutionary history of life during geological time. Fossils mostly consist of mineralized remains or impressions of organisms, however in exceptional cases remnants or derivatives of ancient biomolecules can be preserved and used, for example, to clarify the phylogenetic affinities of enigmatic fossils^[1,2]^. Several analytical approaches have been developed to study these fossilized organics at the molecular level^[2]^. Fossilized pigments in ancient invertebrates, feathered dinosaurs, and mammals were identified by gas chromatography-mass spectrometry (GC-MS) and time of flight secondary ion mass spectroscopy (ToF SIMS), and used to infer the original colours and colour patterns of these extinct organisms^[3]^. The identification of chitin and/or chitosan by Fourier-transform infrared (FTIR) spectroscopy in 1-billion-year-old microfossils, in conjunction with detailed morphological and ultrastructural features documented by transmission electron microscopy (TEM), was used to interpret them as the earliest fungi^[4]^. Taking advantage of some of the previously mentioned tools, as well as other advanced spectroscopic techniques such as scanning transmission X-ray microscopy (STXM) coupled with X-ray absorption near-edge structure (XANES) spectroscopy, numerous studies have highlighted that organic (bio)molecules can sometimes experience only partial degradation during diagenesis and even metamorphism, and be identified in the geological record^[5-14]^.

Recently, preservation of diverse organic degradation products of biomolecules in more than a hundred different metazoan fossils was inferred from spectroscopy data collected with a Raman microspectrometer using a 532 nm laser as the excitation source under continuous illumination^[15-20]^. The reported spectroscopic data were interpreted as evidence for the preservation of organic pigments in eumaniraptoran dinosaur eggshells^[15]^ and in a non-avian dinosaur skin^[18]^, as well as of protein, lipid and/or sugar fossilization products in fossil bones^[16]^, dinosaur eggshells^[20]^, and vertebrate and invertebrate soft-tissues^[17,19]^. Unfortunately, the purported claims of biomolecules in these fossils are not well supported by the data provided, which actually result from instrumental and analytical artefacts. In this paper, we outline the limitations of Raman spectroscopy with respect to the identification of biomolecules in fossil materials, and then describe in detail the origin of the misinterpreted signal.

### Raman spectroscopy and its limitations in the study of organic fossils

Raman spectroscopy is widely used in geosciences as it probes the vibration modes of chemical bonds in both solids, liquids, and gases, with minimal sample preparation ^[21]^. Yet, there are several limitations in terms of the sensitivity and accessibility of chemical fingerprints with the technique as used in the studies questioned here. First, excitation with a 532-nm laser only provides specific information on C-C bonds, and not other covalent linkages, in diagenetically altered organic materials such as fossils^[22]^. As a result, Raman spectra of organic materials preserved in (meta)sedimentary rocks display only the so-called graphite (G) and defect (D1-D4) bands, which provide information about the structural organization of the aromatic skeleton^[23]^. Consistently, Raman spectra of geologically altered organic materials can be similar even when they have significantly different elemental and molecular compositions ^[13,14,24-26]^. Second, under continuous illumination, luminescence occurs concurrently with Stokes Raman scattering and generates a signal that overlaps with the Raman spectral window^[21,27]^. Cross sections of Raman (the probability that Raman scattering takes place) are typically 8 to 10 orders of magnitude smaller than that of luminescence. As a result, a number of precautions are often necessary to be able to detect and interpret Raman spectral features among a number of other spectral variations.

### The reported periodic broad bands are not Raman features

In all the studies questioned here^[15-20]^, the spectral bands assigned to organic molecules are broader than the bands usually associated with Raman scattering, and appear quasi-periodic, in contrast to the non-periodic spectral features typically attributed to Raman scattering of organic compounds.

We investigated the periodicity of these bands using wavelet transform (Fig. 1), an effective signal processing technique that is used to decompose a distorted signal into different frequency scales at various resolution levels. Unlike classical Fourier spectral analyses, wavelet transform analyses are advantageous in describing non-stationarities, i.e. localized variations in frequency or magnitude, and providing a direct visualization of the changing statistical properties. It has become a common tool for analysing localized variations within a time series^[28,29]^, but also for spike removal, denoising and background elimination of Raman spectra^[30,31]^. We selected one spectrum from each of the two studies for which data were made available^[15,19]^. For the wavelet analysis displayed in Fig. 1a,b, we selected the spectrum corresponding to the eggshell of the extant flightless bird *Rhea americana*^[15]^, as it is more likely that a pigment is preserved in a modern sample rather than in a fossil. For the wavelet analysis displayed in Fig. 1c,d, we selected the spectrum collected from the crustacean *Acanthotelson stimpsoni* specimen YPM52348^[19]^, as the chitin–protein complex of crustacean cuticles has a high preservation potential^[8,32]^, and this specimen appears to be one of the best preserved (see fig. 1f in ^[19]^), with the spectrum clearly measured from the specimen (unlike for the specimen shown in fig. 1d of ^[19]^). Note that these two spectra, as well as all other reported ones, were provided by the original authors as baseline-subtracted spectra, not as raw data.

**Figure 1.**
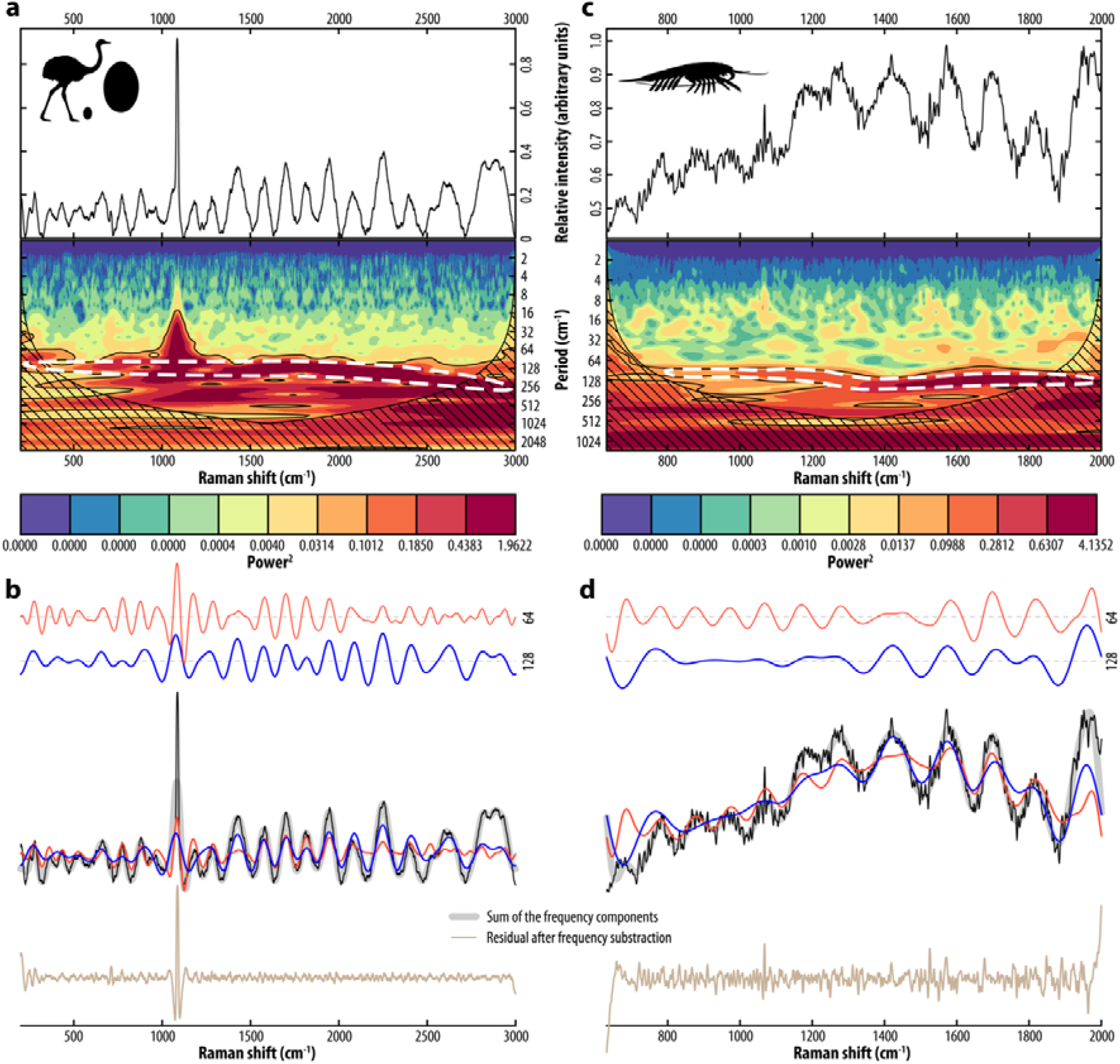
Periodic wavelet analysis of Raman spectra from the eggshell of the extant flightless bird *Rhea americana* (a,b; data from ^[15]^), and from the Carboniferous crustacean *Acanthotelson stimpsoni* specimen YPM52348 (c,d; data from ^[19]^). The hatched area marks parts of the spectrum where energy bands are likely to appear less powerful than they actually are. **a**,**c)** Baseline-subtracted spectra and their wavelet transform analysis show a clear high-frequency periodicity of 64–128 cm^-1^. **b**,**d)** 64 and 128 cm^-1^ frequency components extracted from a wavelet multiresolution analysis (top, in red and blue, respectively) and superimposed, together with the sum of all frequency components, on the spectra. For clarity, the residuals after frequency subtraction are shifted down along the vertical axis.

Both spectra display numerous broad bands for which our wavelet transform analysis reveals clear high-frequency periodicities of ∼64-96 cm^-1^ for wavenumber shifts <1000–1200 cm^-1^, and of ∼128 cm^-1^ for higher wavenumber shifts (Fig. 1a,c). Similar high-frequencies of 130.9 cm^-1^ are obtained by Fast Fourier Transform. The 1086 cm^-1^ carbonate Raman peak present in the *R. americana* spectrum reflects the calcified composition of the eggshell, in contrast to all the other (broader) bands, which are best described as a superposition of quasi-periodic wavelets (Fig. 1b,d). These broad, quasi-periodic bands are not the consequence of any Raman effect, but rather result from physical and instrumental artefacts. Indeed, when a sample is illuminated by the laser, the presence of structural defects and inorganic/organic components can generate significant luminescence, often overwhelming the weak Raman signal^[21,27]^. When this background luminescence is intense, the transmission properties of the interferometric edge filter used to reject the Rayleigh line induce quasi-periodic “ripples” in the measured spectrum^[33]^.

To further illustrate this point, we performed a wavelet analysis on a transmission spectrum of a 532 nm RazorEdge^®^ ultrasteep long-pass edge filter, provided by the manufacturer (Semrock), that is designed to be used as an ultrawide and low-ripple passband edge filter for Raman spectroscopy (Fig. 2). The transmission spectrum of the filter exhibits the aforementioned ripples (Fig. 2a,b). Our wavelet analysis highlights high-frequency periodicities of 64-96 cm^-1^ for low wavenumbers, and of 128 cm^-1^ for higher wavenumbers (Fig. 2b,c), similar to the results reported in the studies questioned herein^[15-20]^. Such edge filter-related instrumental artefacts actually explain the presence of most, if not all, of the broad bands that were attributed to organic molecules.

**Figure 2.**
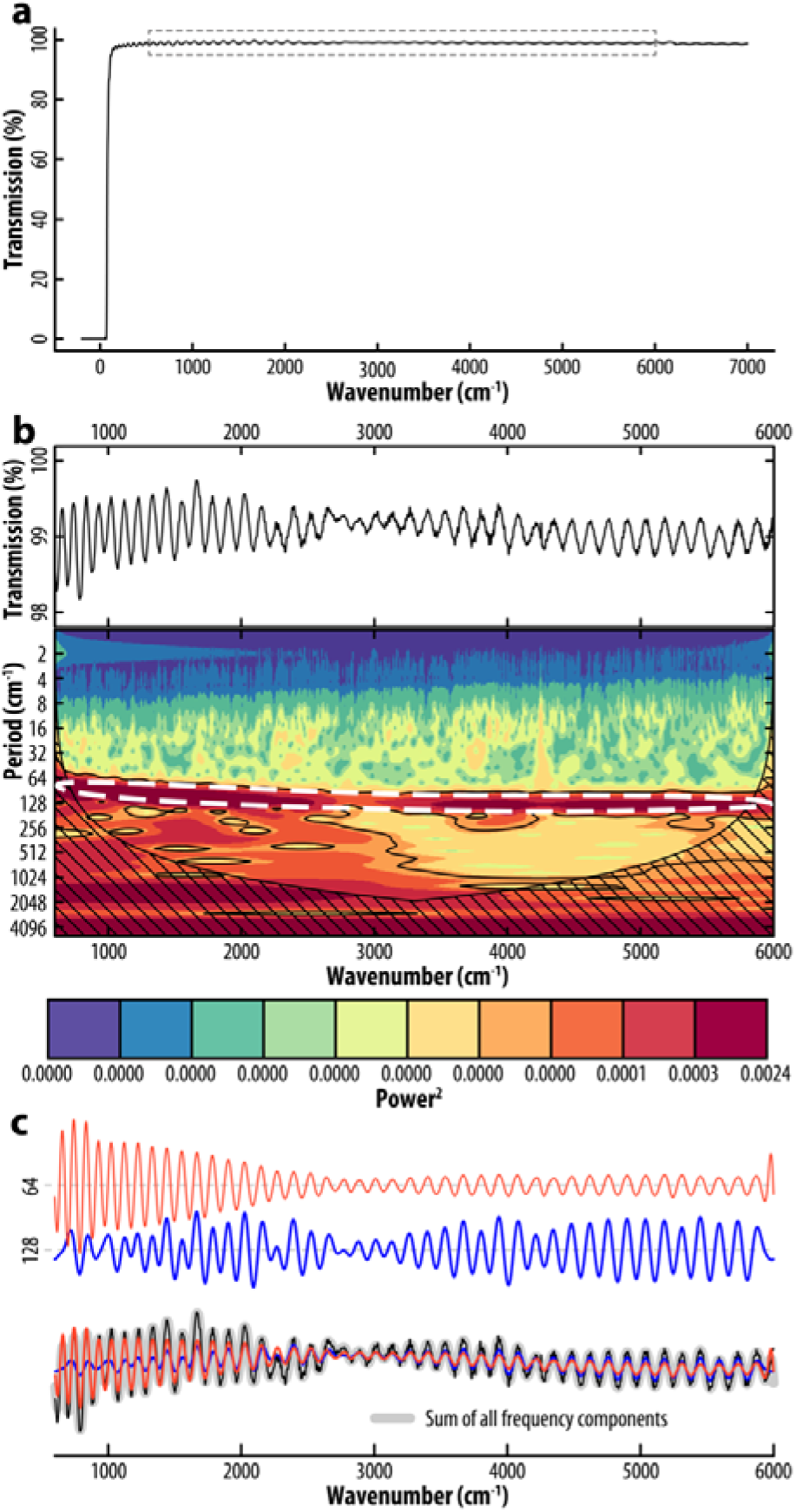
Wavelet transform analysis of the transmission spectrum of a 532 nm RazorEdge^®^ ultrasteep long-pass edge filter (Semrock). **a)** Transmission spectrum of the edge filter between -200 and 7000 cm^-1^. **b)** Wavelet transform analysis of the spectrum between 600 and 6000 cm^-1^ (rectangle in a) showing a clear high-frequency periodicity of 64–128 cm^-1^. **c)** 64 and 128 cm^-1^ frequency components extracted from a wavelet multiresolution analysis (top, in red and blue, respectively) and superimposed, together with the sum of all frequency components, on the spectrum.

### Sample composition does not affect the position of ripples but impacts the shape of the background

The transmission properties of the edge filter induce ripples on the measured spectra when luminescence is intense, making it challenging to identify Raman features without appropriate data processing for background subtraction^[33]^. The data provided in the publications questioned here^[15-20]^ are only the baseline-subtracted spectra, not the raw data, which makes it impossible to precisely assess the impact of non-Raman processes and sample composition on the corrected spectra from which the presence of organic molecules was inferred. To address these issues, we collected Raman microspectroscopy data on modern and fossil crustaceans in analytical conditions similar to those of the aforementioned studies (for details, see Material and Methods in SI).

We reproduced the experiment performed by McCoy et al.^[19]^ using a specimen of the crustacean *Peachocaris strongi* (Fig. 3a) from the same fossil locality (Mazon Creek, Carboniferous, USA). As with other fossils from Mazon Creek, this specimen is preserved as aluminosilicates and calcite in a sideritic concretion (Fig. S1). In order to further assess the impact of the sample’s chemical composition on the measured spectra, we also performed Raman spectroscopy on (i) a specimen of the penaeid shrimp *Cretapenaeus berberus* from the Cretaceous of Morocco (Fig. 3b) preserved as a mixture of calcium phosphates and iron oxides in an illite mudstone (Fig. S1; see also Gueriau et al.^[34]^ and references therein), and (ii) a specimen of the modern shrimp *Neocaridina davidii* (Fig. 3c) dried after death and still rich in organic carbon, likely in the form of chitin (Fig. S1). Whether or not organic carbon is present, and whatever the mineralogical composition of the specimen or its mineral matrix, all the measured spectra (Fig. 3d, solid lines) display broad bands, which all occur at the same wavenumber shifts and add up to a significant background (Fig. 3d, dotted lines). Yet, the shape of the background differs significantly from one measurement to another, and the more intense it is, the more the ripples are expressed. In the baseline-subtracted spectra, the differences in the relative intensity between bands from one measurement to another (Fig. 3e) only result from distinct background profiles of the measurements. A wavelet transform analysis reveals high-frequency periodicities of 64–128 cm^-1^ (Fig. 3f), as was the case for the spectra questioned in the previous section^[15-20]^. Finally, other than the presence of sharp peaks around 964 and 1086 cm^-1^ (Raman peaks of fluorapatite and calcite, respectively), as well as one unidentified peak at 1156 cm^-1^ in the modern shrimp, spectral differences are limited to relative variations in the ripple band intensity that result from analytical artefacts.

**Figure 3.**
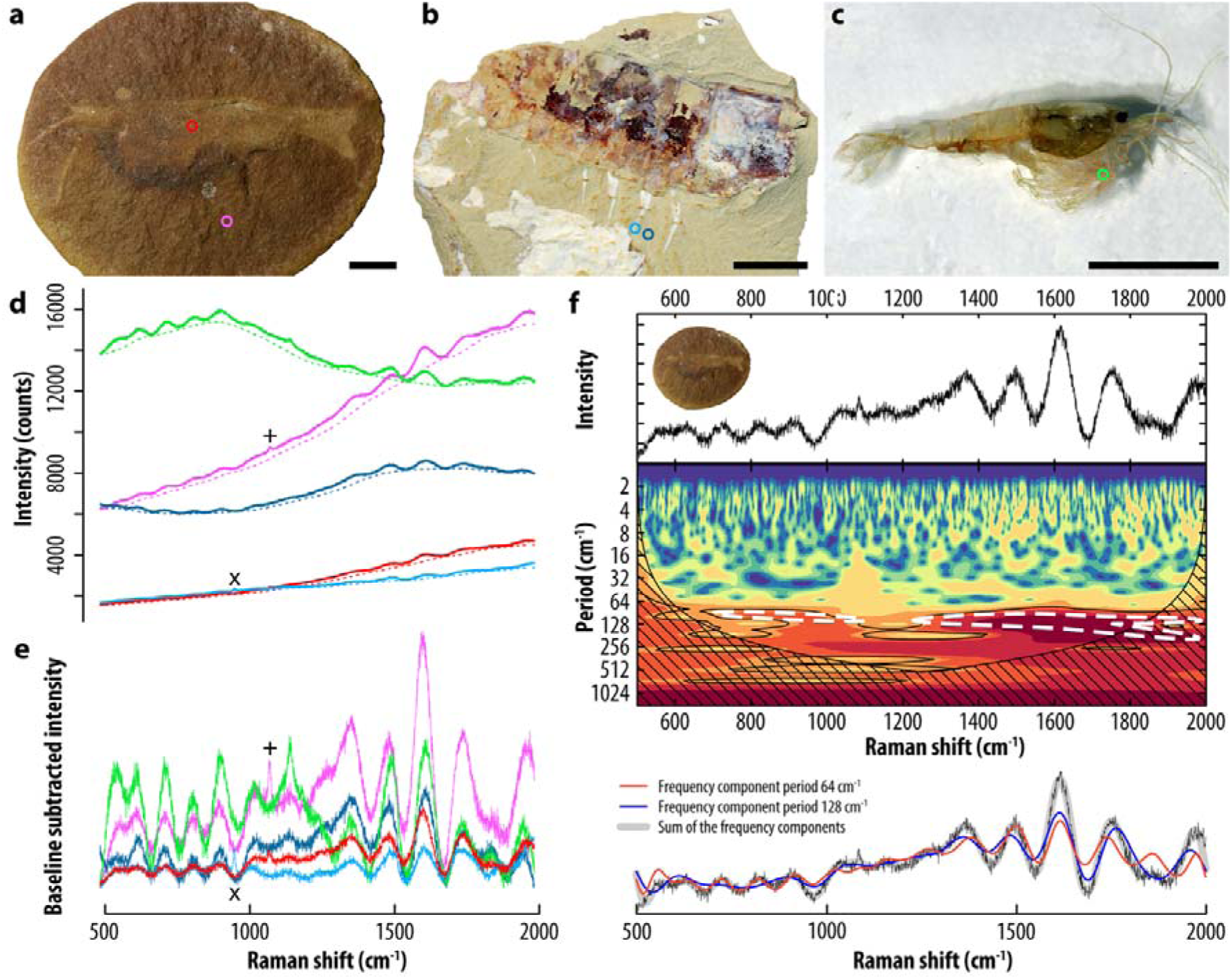
Raman spectroscopic data of fossil and modern shrimps of different chemistry. **a–c)** Photographs of the Carboniferous shrimp *Peachocaris strongi* from Mazon Creek, USA [specimen MGL.107330] (a), the Cretaceous penaeid shrimp *Cretapenaeus berberus* from OT1, Morocco [specimen MHNM-KK-OT 52a] (b), and the extant shrimp *Neocaridina davidii* dried (c). **d)** Raw spectra collected from the areas identified by circles in a–c (solid line), and their baseline (dotted line) as modeled in Spectragryph 1.2 using a 15% adaptive baseline; **e)** corresponding baseline-subtracted spectra. Nearly all bands account for instrumental artefact due to non-linear transmission of the edge filter. Only the sharp peaks highlighted by × and + around 964 and 1086 cm^-1^ (fluorapatite and calcite peaks, respectively) in d and e represent Raman signal. **f)** Wavelet transform analysis of the spectrum collected from *P. strongi* (red in e) showing a high-frequency periodicity between 64 and 128 cm^-1^. Scale bars represent 5 mm.

In short, the ripples observed in the Raman microspectroscopy data questioned here represent remnant instrumental signals that result from confounding broad luminescence and inappropriate data processing. The broad luminescence transmitted by the edge filter induced the ripple-shape features above the cut-off wavelength on the raw spectrum. Background correction did not eliminate the ripple-shape distortions induced, and instead accentuated them to give the appearance of putative signatures of organic molecules.

## Conclusion and Outlook

Broad bands interpreted to be Raman signatures of diverse organic molecule degradation products in various metazoan fossils^[15-20]^ are artefactual quasi-periodic ripples induced by the edge filter due to intense luminescence, and there is no evidence for any preserved organic molecular information. Raman microspectroscopy as used in these papers does not provide information about fossil biomolecules^[22]^, but rather informs on the degree of crystallization of carbonaceous materials, which can be used to quantify the peak temperature they reached during geological burial^[23]^.

Raman spectroscopy can thus provide crucial information about the chemical structure of organic materials, but requires robust and optimized analytical strategies and/or data processing. Several methods have been developed to remove, *a posteriori*, the undesired contribution of luminescence and ripples in Raman spectra^[33,35]^. Note that in the papers discussed here^[15-20]^, such processing would leave no signal other than the mineral peaks. Alternatively, non-conventional time-resolved Raman spectroscopy offers new ways to limit or exploit luminescence signals, while techniques based on coherent anti-Stokes Raman scattering (CARS), surface-enhanced Raman spectroscopy (SERS), and ultraviolet resonance Raman spectroscopy, allow the Raman signal to be considerably enhanced (see Beyssac^[27]^ for review). Furthermore, synchrotron-based X-ray Raman scattering can probe the chemical speciation of light elements such as carbon, in heterogeneous materials usually encountered in life, earth, environmental and materials sciences^[36,37]^.

The search for biomolecules in fossils is a very exciting field of research, offering critical knowledge on both evolutionary events and fossilization processes, yet conventional Raman spectroscopy cannot be used to identify fossil biomolecules. Instead, non-conventional Raman spectroscopy, mass spectrometry and infrared and X-ray absorption spectroscopy techniques, allow paleontologists to correctly identify fossil biomolecules in the geological record^[2,38]^.

## Material and Methods

### Materials used for the different analyses

In Fig. 1, we reused already published baseline-subtracted spectra provided along the original publications by Wiemann et al.^[15]^ and McCoy et al.^[19]^. Specifically, we used the spectrum collected from the eggshell of the extant flightless bird *Rhea americana* ^[15]^, and the spectrum collected from the fossil crustacean *Acanthotelson stimpsoni* specimen YPM52348 from the Carboniferous (ca. 307 Ma) Mazon Creek Lagerstätte, USA ^[19]^.

The transmission spectrum of the 532 nm RazorEdge^®^ ultrasteep long-pass edge filter (Semrock) analyzed in Fig. 2 is available online on the manufacturer’s website. URL:*https://www.semrock.com/FilterDetails.aspx?id=LP03-532RE-25.*

Fossil specimens studied in Fig. 3a and 3b come from the Carboniferous (ca. 307 Ma) Mazon Creek Lagerstätte, USA, and the Upper Cretaceous (ca. 95 Ma) Jbel oum Tkout Lagerstätte from southeastern Morocco, and are housed at the Musée cantonal de géologie de Lausanne (MGL; Lausanne, Switzerland), and the Muséum national d’Histoire naturelle (MNHN; Paris, France), respectively. Note, however, that the latter material belongs to the Musée d’Histoire naturelle de Marrakech (MHNM; Marrakech, Morocco) and is currently only housed at the MNHN for study, within an agreement between both museums. Requests for materials should be addressed to Antoine Pictet or Nour-Eddine Jalil for the MGL and MHNM specimens, respectively.

The dried specimen of the modern shrimp *Neocaridina davidii* studied in Fig. 3c come from the aquarium lab of the University of Lausanne. An adult *N. davidii* specimen was used in this study. Live specimens were purchased commercially through aquarium suppliers in Nyon, Switzerland. The shrimp were cultured in freshwater aquaria at the University of Lausanne, Switzerland. The specimen used was euthanized via anaesthesia by overdose of clove oil solution for 10 minutes, and then was washed and rinsed in de-ionised water continuously until all oily residue was removed, and finally left to dry.

### Raman microspectroscopy

Raman data were collected in analytical conditions similar to those described by Wiemann et al. ^[17]^, using a Horiba Jobin Yvon LabRAM 800 HR spectrometer (UNIL, Lausanne, Switzerland) in a confocal configuration, equipped with an Ar^+^ laser (532 nm) excitation source and an electron multiplying charge-coupled device (EMCCD). Measurements were performed at constant room temperature, directly on the sample surface, by focusing the laser beam with a 300 µm confocal hole using a long working distance ×50 objective (NA = 0.70). This configuration provided a ≈ 2 µm spot size for a laser power delivered at the sample surface below 1 mW. Light was dispersed using a 1800 gr/mm diffraction grating.

### Wavelet analyses

Continuous wavelet transform and wavelet multiresolution analyses were performed in R using the *dplR* and *waveslim* packages, respectively. In this study, the Morlet wavelet (a Gaussian-modulated sinewave) was chosen for the continuous wavelet transform, and we used a 95% level for the significance test. The hatched area delineating the edges of the spectrum (so-called “cone of influence”) marks parts of the spectrum where energy bands are likely to appear less powerful than they actually are because of the increasing importance of edge effects.

All data and the R script used in this work are publicly available via the following Dryad Digital Repository: Alleon J, Montagnac G, Reynard B, Brulé T, Thoury M, Gueriau P. 2020. Data from: Pushing Raman spectroscopy over the edge: purported signatures of organic molecules in fossils are instrumental artefacts. Dryad Digital Repository: https://doi.org/10.5061/dryad.280gb5mp0.

### Baseline subtraction

Baseline subtraction (Fig. 3d, dotted line) was performed using the SpectraGryph 1.2 spectroscopic software (adaptive baseline, 15%, no offset, minimally smoothed through rectangular averaging over an interval of 4 points), following protocols of the questioned studies^[15-20]^. Depending on the composition of the sample, the shape and quality of the baseline fit, and therefore the subtracted spectrum (Fig. 3e) varies when using the same snuggling curved baseline percentage for all spectra instead of adapting the parameter to each spectrum.

### Scanning electron microscopy

SEM analyses were performed in variable pressure mode, using a Field Emission Zeiss GeminiSEM 500 (UNIL, Lausanne, Switzerland), on specimens mounted on aluminum stubs without any additional preparation. Analyses were conducted at a 10-kV accelerating voltage, using a BSE detector for imaging and an EDX detector X-max 150 (Oxford Instruments) for energy-dispersive X-ray spectroscopy.

## Supporting information

All spectra used in this work

R script

## Acknowledgments

We thank Olivier Reubi (UNIL) for his help with Raman spectroscopy, Louise Jensen (EPFL) for her help with scanning electron microscopy, Orla Bath Enright (UNIL) for providing the euthanized specimen of the modern shrimp *Neocaridina davidii*., and Allison Daley (UNIL) for her English edits and suggestions that helped us improve the clarity of the manuscript. Antoine Pictet (MGL), and Didier Dutheil and Nour-Eddine Jalil (MNHN) provided catalogue numbers for the MGL and MHNM specimens studied herein. This research was conducted in accordance with the University of Lausanne’s ethical policy on the use of animals in experiments. This work is a contribution to the Swiss National Science Foundation project CRSK-2_190580 (PI: P.G.), which funded the research and supported P.G. J.A. was supported by the European Union’s Horizon H2020 research and innovation program ERC (STROMATA, grant agreement 759289; PI: Johanna Marin-Carbonne).

