## Supplementary material for "Pushing Raman spectroscopy over the edge: purported signatures of organic molecules in fossils are instrumental artefacts": Figure S1

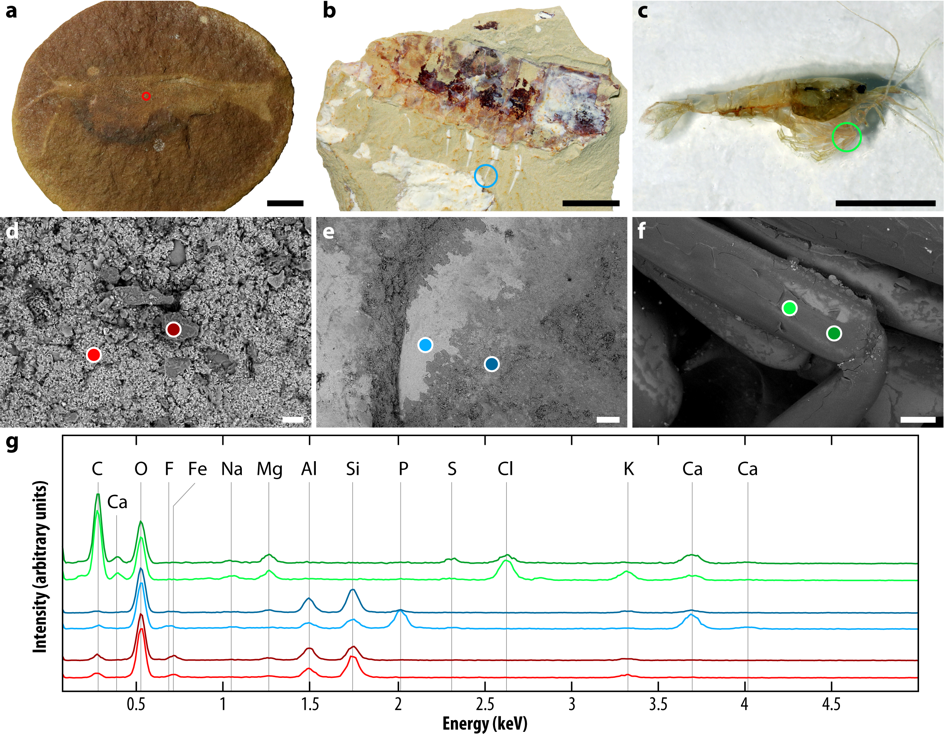

**Figure S1**. Optical **(a–c)** and SEM-BSE **(d–f)** images and corresponding EDX spectra **(g)** of the crustacean specimens investigated here.
