## Supplementary material for "Pushing Raman spectroscopy over the edge: purported signatures of organic molecules in fossils are instrumental artefacts": R script

##########################################################################
###### Load the needed librairies
library(TSA) ## FFT
library(dplR) ## Wavelet analysis
library(waveslim) ## Wavelet Multiresolution analysis (MRA)

###### Load the needed objects and/or functions
specCols <- c("#5E4FA2", "#3288BD", "#66C2A5", "#ABDDA4", "#E6F598", "#FEE08B", "#FDAE61", "#F46D43", "#D53E4F", "#9E0142") # color scale for Wavelet analysis power plots
##########################################################################


###################################################################################
########### Open all the data
###################################################################################

Alleon_etal_data <- read.table("/Users/pgueriau/Dropbox/2-articles-en-cours/BioEssays-Raman/data-processing/Dryad_unified_code_and_data/Alleon_etal_data.txt", skip=2)

###### Fig. 1
Wiemann_etal_RamanShift <- Alleon_etal_data$V1 # cm-1
Wiemann_etal_Rhea <- Alleon_etal_data$V2
McCoy_etal_RamanShift<- Alleon_etal_data$V3[294:length(Alleon_etal_data$V3)] # cm-1 // we cut the starting slope
McCoy_etal_YPM52348 <- Alleon_etal_data$V4[294:length(Alleon_etal_data$V4)] # we cut the starting slope

###### Fig. 2
Filter_wavelength <- Alleon_etal_data$V5 # nm
Filter_transmission <- Alleon_etal_data$V6 # 0-1

###### Fig. 3
Alleon_etal_RamanShift <- Alleon_etal_data$V7
Alleon_etal_Peachocaris_matrix_raw <- Alleon_etal_data$V8
Alleon_etal_Peachocaris_fossil_raw <- Alleon_etal_data$V9
Alleon_etal_Cretapenaeus_matrix_raw <- Alleon_etal_data$V10
Alleon_etal_Cretapenaeus_fossil_raw <- Alleon_etal_data$V11
Alleon_etal_Neocaridina_raw <- Alleon_etal_data$V12
Alleon_etal_Peachocaris_matrix_baseline <- Alleon_etal_data$V13 ## Baseline modeling and subtraction were performed using the SpectraGryph 1.2 spectroscopic software (adaptive baseline, 15%, no offset, minimally smoothed through rectangular averaging over an interval of 4 points), following protocols in (Wiemann et al., 2018a, 2018b, 2020; Fabbri et al., 2020; McCoy et al., 2020; Norell et al., 2020)
Alleon_etal_Peachocaris_matrix_baseline_subtracted <- Alleon_etal_data$V14
Alleon_etal_Peachocaris_fossil_baseline <- Alleon_etal_data$V15
Alleon_etal_Peachocaris_fossil_baseline_subtracted <- Alleon_etal_data$V16
Alleon_etal_Cretapenaeus_matrix_baseline <- Alleon_etal_data$V17
Alleon_etal_Cretapenaeus_matrix_baseline_subtracted <- Alleon_etal_data$V18
Alleon_etal_Cretapenaeus_fossil_baseline <- Alleon_etal_data$V19
Alleon_etal_Cretapenaeus_fossil_baseline_subtracted <- Alleon_etal_data$V20
Alleon_etal_Neocaridina_baseline <- Alleon_etal_data$V21
Alleon_etal_Neocaridina_baseline_subtracted <- Alleon_etal_data$V22


###################################################################################
###################################################################################
############# Fig. 1 - Periodicity data Wiemann et al; McCoy et al ################
###################################################################################
###################################################################################


###################################################################################
###### Fig. 1a,b - Wiemann et al. 2018 Nature
###################################################################################

###### plot the spectra to check that everything is okay
quartz()
plot(x=Wiemann_etal_RamanShift, y=Wiemann_etal_Rhea, t="l", xlab="Raman shift (cm-1)", ylab="Intensity/counts", main="", col="green", ylim=c(0,1))


###### Resample the spectrum to a regularly spaced 1 cm-1 shift
Wiemann_etal_Rhea_spline <-  splinefun(Wiemann_etal_RamanShift, Wiemann_etal_Rhea, method="fmm", ties=mean)
Wiemann_etal_Rhea_spline_1cm1 <- Wiemann_etal_Rhea_spline(seq(200,3000,1))


###### plot it along with the original spectrum to check
quartz()
plot(x=seq(200,3000,1), y=Wiemann_etal_Rhea_spline_1cm1, t="l", xlab="Raman shift (cm-1)", ylab="Transmission (%) / Intensity/counts", main="")
lines(x=Wiemann_etal_RamanShift, y=(Wiemann_etal_Rhea), col="green")


############
######  FFT
############

###### periodicity library(TSA)
#### plot the periodogram
quartz()
p_Wiemann_etal_Rhea_1cm1 = periodogram(Wiemann_etal_Rhea_spline_1cm1, xlim=c(0,0.02))
#### The periodogram shows the “power” of each possible frequency, and we can clearly see spikes at around frequency 0.15 Hz

#### show values for the highest 8 spikes
dd_Wiemann_etal_Rhea_1cm1 = data.frame(freq=p_Wiemann_etal_Rhea_1cm1$freq, spec=p_Wiemann_etal_Rhea_1cm1$spec)
order_Wiemann_etal_Rhea_1cm1 = dd_Wiemann_etal_Rhea_1cm1[order(-dd_Wiemann_etal_Rhea_1cm1$spec),]
top8_Wiemann_etal_Rhea_1cm1 = head(order_Wiemann_etal_Rhea_1cm1, 8)

#### display the 8 highest "power" frequencies
top8_Wiemann_etal_Rhea_1cm1

##### turn in into time equivalent (1 index = 1 one cm-1 here)
time_Wiemann_etal_Rhea_1cm1 = 1/top8_Wiemann_etal_Rhea_1cm1$f
time_Wiemann_etal_Rhea_1cm1


###########
###### Fig. 1a - Wavelet transform (WT) analysis
###########

Morlet_Wiemann_etal_Rhea_1cm1 <- morlet(y1 = Wiemann_etal_Rhea_spline_1cm1, x1 = seq(200,3000,1), dj = 0.1, siglvl = 0.95)

#### plot the wavelet analysis results
quartz()
wavelet.plot(Morlet_Wiemann_etal_Rhea_1cm1,useRaster = TRUE, key.cols=specCols, x.lab="Raman shift (cm-1)", reverse.y = TRUE)


###########
###### Fig. 1b top - Multi-resolution analysis (MRA) decomposition of the signal
###########

###### MRA
nShift <- length(seq(200,3000,1))
nPwrs2.b <- trunc(log(nShift)/log(2)) - 1

mra_Wiemann_etal_Rhea_1cm1 <- mra(Wiemann_etal_Rhea_spline_1cm1, wf = "la8", J = nPwrs2.b, method = "modwt", boundary = "periodic")
ShiftLabels <- paste(2^(1:nPwrs2.b),".",sep="") # '.' = Raman Shift (cm-1)

quartz()
par(mar=c(3,2,2,2),mgp=c(1.25,0.25,0),tcl=0.5, xaxs="i",yaxs="i")
plot(seq(200,3000,1),rep(1,nShift),type="n", axes=FALSE, ylab="",xlab="", ylim=c(-3,38))
title(main="Multiresolution decomposition of Wiemann_etal_Rhea",line=0.75)
axis(side=1)
mtext("Raman shift (cm-1)",side=1, line = 1.25)
Offset <- 0
mra_Wiemann_etal_Rhea_1cm1_2 <- scale(as.data.frame(mra_Wiemann_etal_Rhea_1cm1))
for(i in nPwrs2.b:1){
    x <- scale(mra_Wiemann_etal_Rhea_1cm1[[i]]) + Offset
    # x <- mra_Wiemann_etal_Rhea_1cm1_2[,i]
    Offset
    lines(seq(200,3000,1),x)
    abline(h=Offset,lty="dashed")
    mtext(names(mra_Wiemann_etal_Rhea_1cm1)[[i]],side=2,at=Offset,line = 0)
    mtext(ShiftLabels[i],side=4,at=Offset,line = 0)
    Offset <- Offset+4
}
box()


###########
###### Fig. 1b bottom - MRA surperposed on the spectrum
###########

###### plot frequency components 64 and 128 cm-1 on top of the spectrum
quartz()
plot(x=seq(200,3000,1), y=Wiemann_etal_Rhea_spline_1cm1, t="l", xlab="Raman shift (cm-1)", ylab="Intensoty/counts", main="", ylim=c(-0.2,1))
lines(x=seq(200,3000,1), y=mra_Wiemann_etal_Rhea_1cm1$D6+mra_Wiemann_etal_Rhea_1cm1$S10, col ="red", lwd=3) #  64 cm-1 period
lines(x=seq(200,3000,1), y=mra_Wiemann_etal_Rhea_1cm1$D7+mra_Wiemann_etal_Rhea_1cm1$S10, col ="blue", lwd=3) #  128 cm-1 period

#### add the sum of all
Wiemann_etal_Rhea_MRAsum <- mra_Wiemann_etal_Rhea_1cm1$D10 + mra_Wiemann_etal_Rhea_1cm1$D9 + mra_Wiemann_etal_Rhea_1cm1$D8 + mra_Wiemann_etal_Rhea_1cm1$D7 +mra_Wiemann_etal_Rhea_1cm1$D6 + mra_Wiemann_etal_Rhea_1cm1$S10
lines(x=seq(200,3000,1), y=Wiemann_etal_Rhea_MRAsum, col ="grey", lwd=3)

#### add the residual
Wiemann_etal_Rhea_MRAresidual <- Wiemann_etal_Rhea_spline_1cm1 - Wiemann_etal_Rhea_MRAsum
lines(x=seq(200,3000,1), y=Wiemann_etal_Rhea_MRAresidual, col ="green", lwd=3)


###################################################################################
##### Fig. 1c,d - McCoy et al. 2020 Geobiology
###################################################################################

###### plot the spectra to check that everything is okay
quartz()
plot(x=McCoy_etal_RamanShift, y=McCoy_etal_YPM52348, t="l", xlab="Raman shift (cm-1)", ylab="Intensity/counts", main="", col="darkgreen")


###### Resample the spectrum to a regularly spaced 1 cm-1 shift
McCoy_etal_YPM52348_spline <-  splinefun(McCoy_etal_RamanShift, McCoy_etal_YPM52348, method="fmm", ties=mean)
McCoy_etal_YPM52348_spline_1cm1 <- McCoy_etal_YPM52348_spline(seq(631,2000,1))


###### plot it along with the original spectrum
quartz()
plot(x=seq(631,2000,1), y=McCoy_etal_YPM52348_spline_1cm1, t="l", xlab="Raman shift (cm-1)", ylab="Transmission (%) / Intensity/counts", main="")
lines(x=McCoy_etal_RamanShift, y=McCoy_etal_YPM52348, col="green")


############
######  FFT
############

###### periodicity library(TSA)
#### plot the periodogram
quartz()
p_McCoy_etal_YPM52348_1cm1 = periodogram(McCoy_etal_YPM52348_spline_1cm1, xlim=c(0,0.02))
#### The periodogram shows the “power” of each possible frequency, and we can clearly see spikes at around frequency 0.15 Hz

#### show values for the highest 8 spikes
dd_McCoy_etal_YPM52348_1cm1 = data.frame(freq=p_McCoy_etal_YPM52348_1cm1$freq, spec=p_McCoy_etal_YPM52348_1cm1$spec)
order_McCoy_etal_YPM52348_1cm1 = dd_McCoy_etal_YPM52348_1cm1[order(-dd_McCoy_etal_YPM52348_1cm1$spec),]
top8_McCoy_etal_YPM52348_1cm1 = head(order_McCoy_etal_YPM52348_1cm1, 8)

### display the 8 highest "power" frequencies
top8_McCoy_etal_YPM52348_1cm1

###### turn in into time equivalent (1 index = 1 one cm-1 here after spline)
time_McCoy_etal_YPM52348_1cm1 = 1/top8_McCoy_etal_YPM52348_1cm1$f
time_McCoy_etal_YPM52348_1cm1


###########
###### Fig. 1c - Wavelet transform (WT) analysis
###########

Morlet_McCoy_etal_YPM52348_1cm1 <- morlet(y1 = McCoy_etal_YPM52348_spline_1cm1, x1 = seq(631,2000,1), dj = 0.1, siglvl = 0.95)

#### plot the wavelet analysis results
quartz()
wavelet.plot(Morlet_McCoy_etal_YPM52348_1cm1,useRaster = TRUE, key.cols=specCols, x.lab="Raman shift (cm-1)", reverse.y = TRUE)


###########
###### Fig. 1d top - Multi-resolution analysis (MRA) decomposition of the signal
###########

###### MRA
nShift <- length(seq(631,2000,1))
nPwrs2.b <- trunc(log(nShift)/log(2))-1

mra_McCoy_etal_YPM52348_1cm1 <- mra(McCoy_etal_YPM52348_spline_1cm1, wf = "la8", J = nPwrs2.b, method = "modwt", boundary = "periodic")
ShiftLabels <- paste(2^(1:nPwrs2.b),".",sep="") # '.' = Raman Shift (cm-1)

quartz()
par(mar=c(3,2,2,2),mgp=c(1.25,0.25,0),tcl=0.5, xaxs="i",yaxs="i")
plot(seq(631,2000,1),rep(1,nShift),type="n", axes=FALSE, ylab="",xlab="", ylim=c(-3,38))
title(main="Multiresolution decomposition of McCoy_etal_YPM52348",line=0.75)
axis(side=1)
mtext("Raman shift (cm-1)",side=1, line = 1.25)
Offset <- 0
mra_McCoy_etal_YPM52348_1cm1_2 <- scale(as.data.frame(mra_McCoy_etal_YPM52348_1cm1))
for(i in nPwrs2.b:1){
    x <- scale(mra_McCoy_etal_YPM52348_1cm1[[i]]) + Offset
    # x <- mra_McCoy_etal_YPM52348_1cm1_2[,i]
    Offset
    lines(seq(631,2000,1),x)
    abline(h=Offset,lty="dashed")
    mtext(names(mra_McCoy_etal_YPM52348_1cm1)[[i]],side=2,at=Offset,line = 0)
    mtext(ShiftLabels[i],side=4,at=Offset,line = 0)
    Offset <- Offset+4
}
box()


###########
###### Fig. 1d bottom - MRA surperposed on the spectrum
###########

###### plot frequency components 64 and 128 cm-1 on top of the spectrum
quartz()
plot(x=seq(631,2000,1), y=McCoy_etal_YPM52348_spline_1cm1, t="l", xlab="Raman shift (cm-1)", ylab="Intensity/counts", main="", ylim=c(-0.1,1))
lines(x=seq(631,2000,1), y=mra_McCoy_etal_YPM52348_1cm1$D6+mra_McCoy_etal_YPM52348_1cm1$S9, col ="red", lwd=3) #  64 cm-1 period
lines(x=seq(631,2000,1), y=mra_McCoy_etal_YPM52348_1cm1$D7+mra_McCoy_etal_YPM52348_1cm1$S9, col ="blue", lwd=3) #  128 cm-1 period

#### add the sum of all
McCoy_etal_YPM52348_MRAsum <- mra_McCoy_etal_YPM52348_1cm1$D9 + mra_McCoy_etal_YPM52348_1cm1$D8 + mra_McCoy_etal_YPM52348_1cm1$D7 + mra_McCoy_etal_YPM52348_1cm1$D6 + mra_McCoy_etal_YPM52348_1cm1$S9
lines(x=seq(631,2000,1), y=McCoy_etal_YPM52348_MRAsum, col ="grey", lwd=3)


#### add the residual
McCoy_etal_YPM52348_MRAresidual <- McCoy_etal_YPM52348_spline_1cm1 - McCoy_etal_YPM52348_MRAsum
lines(x=seq(631,2000,1), y=McCoy_etal_YPM52348_MRAresidual, col ="green", lwd=3)


###################################################################################
###################################################################################
############################## Fig. 2 - Edge filter ###############################
###################################################################################
###################################################################################

###### plot the spectra to check that everything is okay
quartz()
plot(x=Filter_wavelength, y=Filter_transmission*100, t="l", xlab="Wavelength (nm)", ylab="Transmission (%)", main="") # *100 to express the transmission in % instead of 0-1


###### Convert nm to cm-1, considering the use of a 532 nm laser
Filter_wavelength_cm1 <- (1E7/532)-(1E7/Filter_wavelength)


###### plot to check
quartz()
plot(x=Filter_wavelength_cm1, y=Filter_transmission*100, t="l", xlab="Wavelength (cm-1)", ylab="Transmission (%)", main="") # use xlim=c(0,3000)) to focus on the Raman wavelength range


###### Resample the spectrum to a regularly spaced 1 cm-1 shift
Filter_transmission_spline <-  splinefun(Filter_wavelength_cm1, Filter_transmission, method="fmm", ties=mean)
Filter_transmission_spline_1cm1 <- Filter_transmission_spline(seq(-200,7000,1))


###### plot it along with the original spectrum
quartz()
plot(x=Filter_wavelength_cm1, y=Filter_transmission*100, t="l", xlab="Wavelength (cm-1)", ylab="Transmission (%)", main="", xlim=c(-200,7000))
lines(x=seq(-200,7000,1), y=Filter_transmission_spline_1cm1*100, col="green")


###################################################################################
###### Fig. 2a - Edge filter transmission
###################################################################################

quartz()
plot(x=seq(-200,7000,1), y=Filter_transmission_spline_1cm1*100, t="l", xlab="Wavelength (cm-1)", ylab="Transmission (%)", main="")


###################################################################################
###### Fig. 2b,c - Edge filter periodicty
###################################################################################

###### Re-Resample the spectrum to the oscillatory range
Filter_transmission_spline_1cm1_osc <- Filter_transmission_spline(seq(600,6000,1))

###### plot it to check
quartz()
plot(x=seq(600,6000,1), y=Filter_transmission_spline_1cm1_osc*100, t="l", xlab="Wavenumber (cm-1)", ylab="Transmission (%)", main="")


############
######  FFT
############

###### periodicity library(TSA)
#### plot the periodogram
quartz()
p_filter = periodogram(Filter_transmission_spline_1cm1_osc, xlim=c(0,0.02))

#### show values for the highest 8 spikes
dd_filter = data.frame(freq=p_filter$freq, spec=p_filter$spec)
order_filter = dd_filter[order(-dd_filter$spec),]
top8_filter = head(order_filter, 8)

### display the 8 highest "power" frequencies
top8_filter

##### turn in into time equivalent (1 index = 1 one day here)
filterStep <- 1 # cm-1 after spine
time_filter = filterStep/top8_filter$f
time_filter


############
###### Fig. 2b - Wavelet transform (WT) analysis
############

Morlet_filter_1cm1 <- morlet(y1 = Filter_transmission_spline_1cm1_osc, x1 = seq(600,6000,1), dj = 0.1, siglvl = 0.95)

#### plot the wavelet analysis results
quartz()
wavelet.plot(Morlet_filter_1cm1,useRaster = TRUE, key.cols=specCols, x.lab="Wavenumber (cm-1)", reverse.y = TRUE, crn.ylim = c(0.98,1.0))


############
###### Fig. 2c top - Multi-resolution analysis (MRA) decomposition of the signal
############

###### MRA
nShift <- length(seq(600,6000,1))
nPwrs2.b <- trunc(log(nShift)/log(2))-1

mra_filter_1cm1 <- mra(Filter_transmission_spline_1cm1_osc, wf = "la8", J = nPwrs2.b, method = "modwt", boundary = "periodic")
ShiftLabels <- paste(2^(1:nPwrs2.b),".",sep="") # '.' = Raman Shift (cm-1)

quartz()
par(mar=c(3,2,2,2),mgp=c(1.25,0.25,0),tcl=0.5, xaxs="i",yaxs="i")
plot(seq(600,6000,1),rep(1,nShift),type="n", axes=FALSE, ylab="",xlab="", ylim=c(-3,38))
title(main="Multiresolution decomposition of the edge filter",line=0.75)
axis(side=1)
mtext("Wavenumber (cm-1)",side=1, line = 1.25)
Offset <- 0
mra_filter_1cm1_2 <- scale(as.data.frame(mra_filter_1cm1))
for(i in nPwrs2.b:1){
    x <- scale(mra_filter_1cm1[[i]]) + Offset
    # x <- mra_filter_1cm1_2[,i]
    Offset
    lines(seq(600,6000,1),x)
    abline(h=Offset,lty="dashed")
    mtext(names(mra_filter_1cm1)[[i]],side=2,at=Offset,line = 0)
    mtext(ShiftLabels[i],side=4,at=Offset,line = 0)
    Offset <- Offset+4
}
box()


###########
##### Fig. 2c bottom - MRA surposed on the spectrum
###########

###### plot frequency components 64 and 128 cm-1 on top of the spectrum
quartz()
plot(x=seq(600,6000,1), y=Filter_transmission_spline_1cm1_osc*100, t="l", xlab="Wavenumber (cm-1)", ylab="Transmission (%)", main="", ylim= c(98,100))
lines(x=seq(600,6000,1), y=(mra_filter_1cm1$D6+mra_filter_1cm1$S11)*100, col ="red", lwd=3) #  64 cm-1 period
lines(x=seq(600,6000,1), y=(mra_filter_1cm1$D7+mra_filter_1cm1$S11)*100, col ="blue", lwd=3) #  128 cm-1 period

#### add the sum of all
filter_MRAsum <- mra_filter_1cm1$D11 + mra_filter_1cm1$D10 + mra_filter_1cm1$D9 + mra_filter_1cm1$D8 + mra_filter_1cm1$D7 + mra_filter_1cm1$D6 + mra_filter_1cm1$S11
lines(x=seq(600,6000,1), y=filter_MRAsum*100, col ="grey", lwd=3)


########
###### if we want to add also the residual
quartz()
plot(x=seq(600,6000,1), y=Filter_transmission_spline_1cm1_osc*100, t="l", xlab="Wavenumber (cm-1)", ylab="Transmission (%)", main="", ylim= c(0,100))
lines(x=seq(600,6000,1), y=(mra_filter_1cm1$D6+mra_filter_1cm1$S11)*100, col ="red", lwd=3) #  64 cm-1 period
lines(x=seq(600,6000,1), y=(mra_filter_1cm1$D7+mra_filter_1cm1$S11)*100, col ="blue", lwd=3) #  128 cm-1 period

#### add the sum of all
filter_MRAsum <- mra_filter_1cm1$D11 + mra_filter_1cm1$D10 + mra_filter_1cm1$D9 + mra_filter_1cm1$D8 + mra_filter_1cm1$D7 + mra_filter_1cm1$D6 + mra_filter_1cm1$S11
lines(x=seq(600,6000,1), y=filter_MRAsum*100, col ="grey", lwd=3)

#### add the residual
Wiemann_etal_Rhea_MRAresidual <- Filter_transmission_spline_1cm1_osc - filter_MRAsum
lines(x=seq(600,6000,1), y=Wiemann_etal_Rhea_MRAresidual*100, col ="green", lwd=3)


###### plot the residual alone to better see it
quartz()
plot(x=seq(600,6000,1), y=Wiemann_etal_Rhea_MRAresidual*100, t="l", , xlab="Wavenumber (cm-1)", ylab="Transmission (%)", col ="green", lwd=3)


###################################################################################
###################################################################################
###################### Fig. 3 - Original data Alleon et al. #######################
###################################################################################
###################################################################################


###################################################################################
###### Fig. 3d - Raw spectra and baselines produced using SpectraGryph 1.2
###################################################################################

quartz()
plot(x=Alleon_etal_RamanShift, y=Alleon_etal_Peachocaris_matrix_raw, t="l", xlab="Raman shift (cm-1)", ylab="Intensity/counts", main="", col="pink", ylim=c(2000,16000))
lines(x=Alleon_etal_RamanShift, y=Alleon_etal_Peachocaris_fossil_raw, col="red")
lines(x=Alleon_etal_RamanShift, y=Alleon_etal_Cretapenaeus_matrix_raw, col="darkblue")
lines(x=Alleon_etal_RamanShift, y=Alleon_etal_Cretapenaeus_fossil_raw, col="lightblue")
lines(x=Alleon_etal_RamanShift, y=Alleon_etal_Neocaridina_raw, col="green")

legend(x=1600, y=11500,legend=c("Peachocaris matrix", "Peachocaris fossil", "Cretapenaeus matrix", "Cretapenaeus fossil", "Neocaridina"), text.col=c("pink", "red", "darkblue", "lightblue", "green"), lwd=1.5,lty=1, col=c("pink", "red", "darkblue", "lightblue", "green"), cex=0.65)

###### add the baselines
lines(x=Alleon_etal_RamanShift, y=Alleon_etal_Peachocaris_matrix_baseline, col="pink", lty=3)
lines(x=Alleon_etal_RamanShift, y=Alleon_etal_Peachocaris_fossil_baseline, col="red", lty=3)
lines(x=Alleon_etal_RamanShift, y=Alleon_etal_Cretapenaeus_matrix_baseline, col="darkblue", lty=3)
lines(x=Alleon_etal_RamanShift, y=Alleon_etal_Cretapenaeus_fossil_baseline, col="lightblue", lty=3)
lines(x=Alleon_etal_RamanShift, y=Alleon_etal_Neocaridina_baseline, col="green", lty=3)


###################################################################################
###### Fig. 3e - Baseline subtracted spectra
###################################################################################

quartz()
plot(x=Alleon_etal_RamanShift, y=Alleon_etal_Peachocaris_matrix_baseline_subtracted, t="l", xlab="Raman shift (cm-1)", ylab="Intensity/counts", main="", col="pink")
lines(x=Alleon_etal_RamanShift, y=Alleon_etal_Peachocaris_fossil_baseline_subtracted, col="red")
lines(x=Alleon_etal_RamanShift, y=Alleon_etal_Cretapenaeus_matrix_baseline_subtracted, col="darkblue")
lines(x=Alleon_etal_RamanShift, y=Alleon_etal_Cretapenaeus_fossil_baseline_subtracted, col="lightblue")
lines(x=Alleon_etal_RamanShift, y=Alleon_etal_Neocaridina_baseline_subtracted, col="green")

legend("topleft",legend=c("Peachocaris matrix", "Peachocaris fossil", "Cretapenaeus matrix", "Cretapenaeus fossil", "Neocaridina"), text.col=c("pink", "red", "darkblue", "lightblue", "green"), lwd=1.5,lty=1, col=c("pink", "red", "darkblue", "lightblue", "green"), cex=0.65)


###################################################################################
###### Fig. 3f - Peachocaris (fossil) periodicty
###################################################################################

###### Resample the spectrum to a regularly spaced 1 cm-1 shift
Alleon_etal_Peachocaris_fossil_baseline_subtracted_spline <-  splinefun(Alleon_etal_RamanShift, Alleon_etal_Peachocaris_fossil_baseline_subtracted, method="fmm", ties=mean)
Alleon_etal_Peachocaris_fossil_baseline_subtracted_spline_1cm1 <- Alleon_etal_Peachocaris_fossil_baseline_subtracted_spline(seq(500,2000,1))


###### plot it along with the original spectrum to check
quartz()
plot(x=Alleon_etal_RamanShift, y=Alleon_etal_Peachocaris_fossil_baseline_subtracted, t="l", xlab="Raman shift (cm-1)", ylab="Intensity/counts", main="")
lines(x=seq(500,2000,1), y=Alleon_etal_Peachocaris_fossil_baseline_subtracted_spline_1cm1, col="green")


############
######  FFT
############

###### periodicity library(TSA)
#### plot the periodogram
quartz()
p_Peachocaris_1cm1 = periodogram(Alleon_etal_Peachocaris_fossil_baseline_subtracted_spline_1cm1, xlim=c(0,0.02))
#### The periodogram shows the “power” of each possible frequency, and we can clearly see spikes at around frequency 0.15 Hz

#### show values for the highest 8 spikes
dd_Peachocaris_1cm1 = data.frame(freq=p_Peachocaris_1cm1$freq, spec=p_Peachocaris_1cm1$spec)
order_Peachocaris_1cm1 = dd_Peachocaris_1cm1[order(-dd_Peachocaris_1cm1$spec),]
top8_Peachocaris_1cm1 = head(order_Peachocaris_1cm1, 8)

### display the 8 highest "power" frequencies
top8_Peachocaris_1cm1

##### turn in into time equivalent (1 index = 1 one cm-1 here after spline)
time_Peachocaris_1cm1 = 1/top8_Peachocaris_1cm1$f
time_Peachocaris_1cm1


############
###### Fig. 3f top – Wavelet transform (WT) analysis
############

Morlet_Peachocaris_1cm1 <- morlet(y1 = Alleon_etal_Peachocaris_fossil_baseline_subtracted_spline_1cm1, x1 = seq(500,2000,1), dj = 0.1, siglvl = 0.95)

#### plot the wavelet analysis results
quartz()
wavelet.plot(Morlet_Peachocaris_1cm1,useRaster = TRUE, key.cols=specCols, x.lab="Raman shift (cm-1)", reverse.y = TRUE)


###### Multi-resolution analysis (MRA) decomposition of the signal

nShift <- length(seq(500,2000,1))
nPwrs2.b <- trunc(log(nShift)/log(2)) - 1

mra_Peachocaris_1cm1 <- mra(Alleon_etal_Peachocaris_fossil_baseline_subtracted_spline_1cm1, wf = "la8", J = nPwrs2.b, method = "modwt", boundary = "periodic")
ShiftLabels <- paste(2^(1:nPwrs2.b),".",sep="") # '.' = Raman Shift (cm-1)

quartz()
par(mar=c(3,2,2,2),mgp=c(1.25,0.25,0),tcl=0.5, xaxs="i",yaxs="i")
plot(seq(500,2000,1),rep(1,nShift),type="n", axes=FALSE, ylab="",xlab="", ylim=c(-3,38))
title(main="Multiresolution decomposition of Peachocaris_fossil_baseline_subtracted",line=0.75)
axis(side=1)
mtext("Raman shift (cm-1)",side=1, line = 1.25)
Offset <- 0
mra_Peachocaris_1cm1_2 <- scale(as.data.frame(mra_Peachocaris_1cm1))
for(i in nPwrs2.b:1){
    x <- scale(mra_Peachocaris_1cm1[[i]]) + Offset
    # x <- mra_Peachocaris_1cm1_2[,i]
    Offset
    lines(seq(500,2000,1),x)
    abline(h=Offset,lty="dashed")
    mtext(names(mra_Peachocaris_1cm1)[[i]],side=2,at=Offset,line = 0)
    mtext(ShiftLabels[i],side=4,at=Offset,line = 0)
    Offset <- Offset+4
}
box()


############
###### Fig. 3f bottom - MRA surposed on the spectrum
############

###### plot frequency components 64 and 128 cm-1 on top of the spectrum
quartz()
plot(x=seq(500,2000,1), y=Alleon_etal_Peachocaris_fossil_baseline_subtracted_spline_1cm1, t="l", xlab="Raman shift (cm-1)", ylab="Intensity/counts", main="")
lines(x=seq(500,2000,1), y=mra_Peachocaris_1cm1$D6+mra_Peachocaris_1cm1$S9, col ="red", lwd=3) #  64 cm-1 period
lines(x=seq(500,2000,1), y=mra_Peachocaris_1cm1$D7+mra_Peachocaris_1cm1$S9, col ="blue", lwd=3) #  128 cm-1 period

#### add the sum of all
Peachocaris_MRAsum <- mra_Peachocaris_1cm1$D9 + mra_Peachocaris_1cm1$D8 + mra_Peachocaris_1cm1$D7 +mra_Peachocaris_1cm1$D6 + mra_Peachocaris_1cm1$S9
lines(x=seq(500,2000,1), y=Peachocaris_MRAsum, col ="grey", lwd=3)


########
###### if we want to add also the residual
quartz()
plot(x=seq(500,2000,1), y=Alleon_etal_Peachocaris_fossil_baseline_subtracted_spline_1cm1, t="l", xlab="Raman shift (cm-1)", ylab="Intensity/counts", main="", ylim= c(-50,400))
lines(x=seq(500,2000,1), y=mra_Peachocaris_1cm1$D6+mra_Peachocaris_1cm1$S9, col ="red", lwd=3) #  64 cm-1 period
lines(x=seq(500,2000,1), y=mra_Peachocaris_1cm1$D7+mra_Peachocaris_1cm1$S9, col ="blue", lwd=3) #  128 cm-1 period

#### add the sum of all
Peachocaris_MRAsum <- mra_Peachocaris_1cm1$D9 + mra_Peachocaris_1cm1$D8 + mra_Peachocaris_1cm1$D7 +mra_Peachocaris_1cm1$D6 + mra_Peachocaris_1cm1$S9
lines(x=seq(500,2000,1), y=Peachocaris_MRAsum, col ="grey", lwd=3)


#### add the residual
Peachocaris_MRAresidual <- Alleon_etal_Peachocaris_fossil_baseline_subtracted_spline_1cm1 - Peachocaris_MRAsum
lines(x=seq(500,2000,1), y=Peachocaris_MRAresidual, col ="green", lwd=3)
